# Identification of Expressed Endogenous Retroviral Element Associated Long Terminal Repeats in Devil Facial Tumour Cells

**DOI:** 10.1101/2025.10.06.679394

**Authors:** Anuk Kruawan, Maximilian Stammnitz, Richard Wilson, Sri H. Ramarathinam, Chrissie E. B. Ong, Kirsten A. Fairfax, Amanda L. Patchett, A. Bruce Lyons, Andrew S. Flies

## Abstract

The Tasmanian devil (*Sarcophilus harrisii*) population has undergone a major decline in the wild due to the epidemics of two transmissible cancers known as devil facial tumours (DFT1 and DFT2). A multipronged conservation strategy is in place, but a vaccine that prevents devils from developing devil facial tumour disease would be a major step towards recovering the wild devil population. Critical to an effective DFT vaccine are target antigens. Putative non-coding regions of genomes have been identified as potential tumour-specific antigens for human cancer. These non-coding regions include long terminal repeats of endogenous retroviral elements. We identified 11,193 ERV LTR transcripts that were expressed in DFT1 or DFT2 transcriptomes but not in healthy tissue samples. Using a proteogenomic approach, we identified 33 ERV LTR peptides unique to DFT1 and/or DFT2 immunopeptidomes; four of these were validated in subsequent screens with synthetic peptides. Our study shows the potential for ERV LTRs as novel vaccine targets against DFT1 and DFT2. This method can be applied to other species for the development of cancer vaccine targets that may be shared across tumour types.

**summary:** ERV LTRs are present in the Tasmanian devil genome and devil facial tumour cell transcriptomes and immunopeptidome.

## Introduction

The wild Tasmanian devil population has been reduced by an estimated 80% due to transmissible cancers known as devil facial tumours (DFTs) [1]. Devil facial tumour 1 (DFT1) was first observed in 1996 and is the primary driver of the population decline [2]. DFT2 is an independent transmissible cancer that was first observed in 2014 and has been detected only in southern Tasmania to date [3, 4]. DFT1 and DFT2 appear grossly similar but they are histologically, cytogenetically, and genetically different [3]. While both DFT1 and DFT2 originated from Schwann-like cells, DFT1 and DFT2 have distinct gene expression profiles [5] and genomic mutations [4]. Due the major conservation and animal welfare issues associated with the transmissible cancers, development of an oral bait vaccine that targets tumour cell proteins is a high management priority [6, 7].

Previous vaccination trials used a whole-cell DFT1 vaccine, which should contain the full repertoire of tumour associated antigens (TAAs) and tumour specific antigens (TSAs) [8]. TAAs are antigens that can be detected in a subset of healthy cells, but are upregulated in tumour cells. TSAs are antigens that arise from mutational or non-canonical antigens and are present only in tumour cells [9-11]. The DFT1 vaccine did not protect devils against tumour establishment following live DFT1 cell challenge. However, an immunotherapy of live MHC-I positive DFT1 cells with adjuvants was able to induce striking tumour regressions in three out of five devils [8]. This demonstrates that the devil immune system can be primed to recognise and eliminate tumour cells.

Despite the ongoing transmission of the DFT1 and DFT2 cells for more than three decades and one decade, respectively, the tumour cells have a limited set of protein-coding mutations that could potentially be targeted by vaccines [4]. Non-canonical tumour antigens have been explored as potential targets for cancer immunotherapy [12]. Endogenous retroviral elements (ERVs) have been increasingly reported as a rich source of non-canonical TSAs [13]. ERVs are remnants of ancient retroviral infections that were incorporated into host genomes and passed on from ancestors to offspring in later generations [14]. **Figure 1*a*** shows the integration of retroviral RNA into the host genome to become an ERV. The full-length integrated ERV contains coding and non-coding domains (**Figure 1*b***). The coding domain consists of the following genes: group-specific antigen (*gag*), protease (*pro*), polymerase (*pol*), and envelope (*env*) (**Figure 1*b***) [15]. The 5’ and 3’ ends of these coding domains consist of long terminal repeats (LTRs), which act as a promoter and enhancer (**Figure 1*b***). ERVs can be inserted as a whole intact gene. However, ERVs are commonly mutated or fragmented, and are silenced or tightly regulated in healthy cells [14, 16, 17] (**Figure 1*b***). Parts of ERVs can be inserted into the host gene or inserted as a stand-alone element as well [17] (**Figure 1*b***).

**Figure 1.**
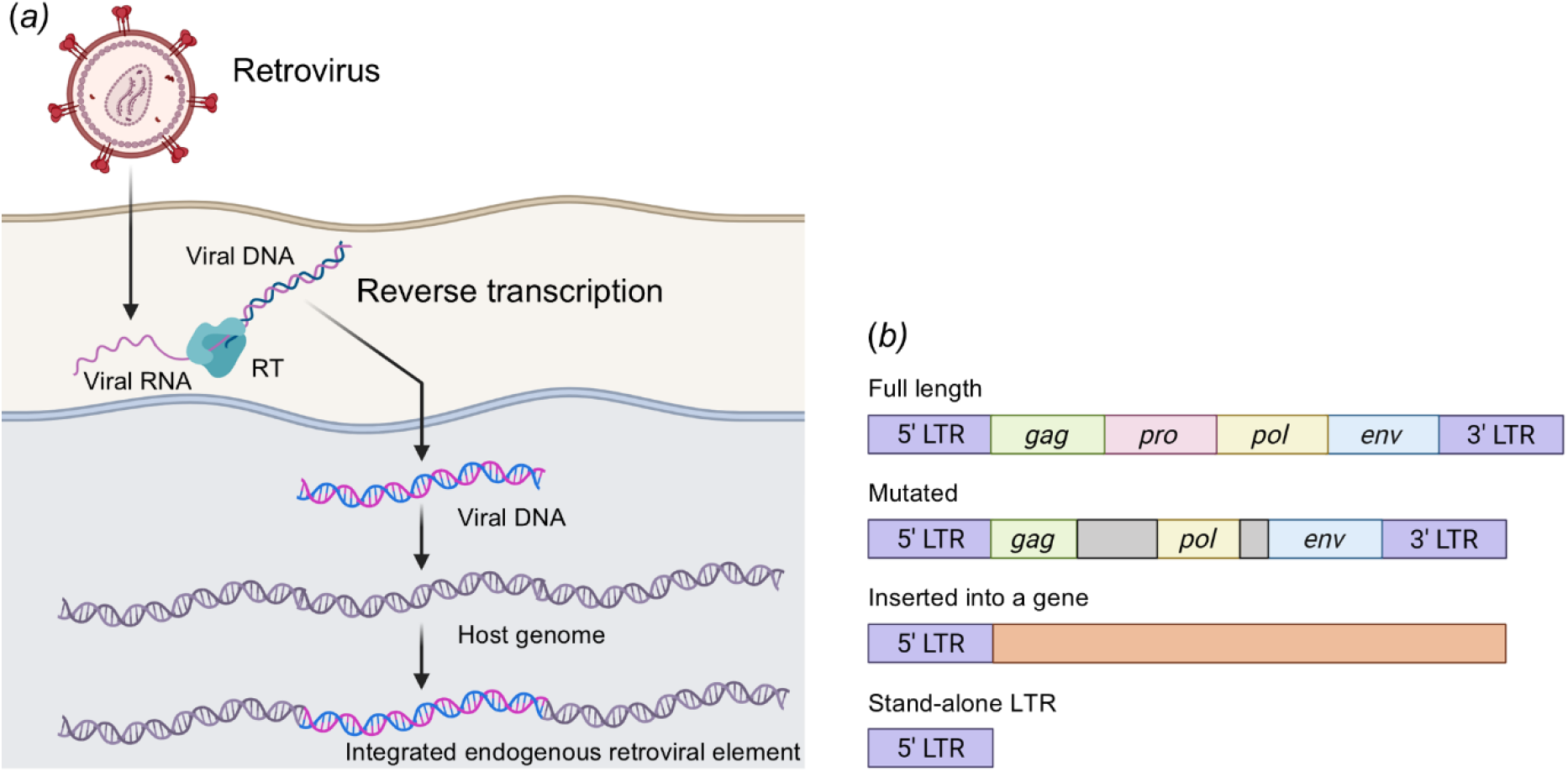
Retroviral DNA integrated into the host cell genome. An illustration showing the integration of retroviral DNA. (*a*) Viral DNA con be integrated as a full-length gene with or without mutation, into a gene found in host genome, or as a stand-alone element. (*b*) Reverse transcriptase (RT), long terminal repeat (LTR), group-specific antigen (*gag*), protease (*pro*), polymerase (*pol*), and envelope (*env*) (created in BioRender, adapted from [17]).

An illustration showing the integration of retroviral DNA. (*a*) Viral DNA con be integrated as a full-length gene with or without mutation, into a gene found in host genome, or as a stand-alone element. (*b*) Reverse transcriptase (RT), long terminal repeat (LTR), group-specific antigen (*gag*), protease (*pro*), polymerase (*pol*), and envelope (*env*) (created in BioRender, adapted from [17]).

The expression of ERVs can produce non-canonical TSAs and the expression of ERVs inserted near genes gene can turn the genes into TAAs through changing their expression pattern [18, 19]. Expression of ERV-derived TAAs and TSAs can be consistent across multiple tumour types, and thus hold great potential for the development of off-the-shelf cancer vaccines [13, 19]. This contrasts with mutationally-derived TSAs, which are generally unique to each cancer.

ERVs and LTRs are present in the Tasmanian devil genome [20-22]. It was reported that the *Dasyuridae* family, which includes Tasmanian devils, have approximately 10 times more ERVs in their genome than other marsupials [22]. Understanding the landscape of ERVs in DFT1 or DFT2 may provide insight into their role in oncogenesis and identify novel TSAs to use in DFT1 and DFT2 vaccines [22]. Here we discovered that thousands of RNA transcripts from ERV LTRs are expressed in DFT1 and DFT2 cells. Furthermore, a subset of these LTRs were detected in DFT1 and/or DFT2 immunopeptidomes, suggesting that a some of transcripts are translated to proteins and presented on major histocompatibility complex class I (MHC-I) proteins for cytotoxic CD8^+^ T cell recognition.

## Materials and Methods

### Overview of the ERV LTR detection pipeline

The ERV LTR detection pipeline comprised of three major sections, namely genomics, transcriptomics, and proteomics (**Figure 2**). First, ERV LTRs were extracted from the Tasmanian devil whole genome sequence (WGS) version mSarHar1.11 [4]. Next, the RNA-seq datasets containing normal (i.e., non-tumour) devil tissue and DFT samples were searched to identify ERV LTR transcripts in DFT cells that were not present in non-DFT samples. DFT-specific ERV LTR transcripts were then six-frame translated to generate a list of peptide sequences, which were then used to create a custom search database for raw mass spectrometry data. The ERV LTR peptides detected in the immunopeptidome were then synthesised for further validation by mass spectrometry.

**Figure 2.**
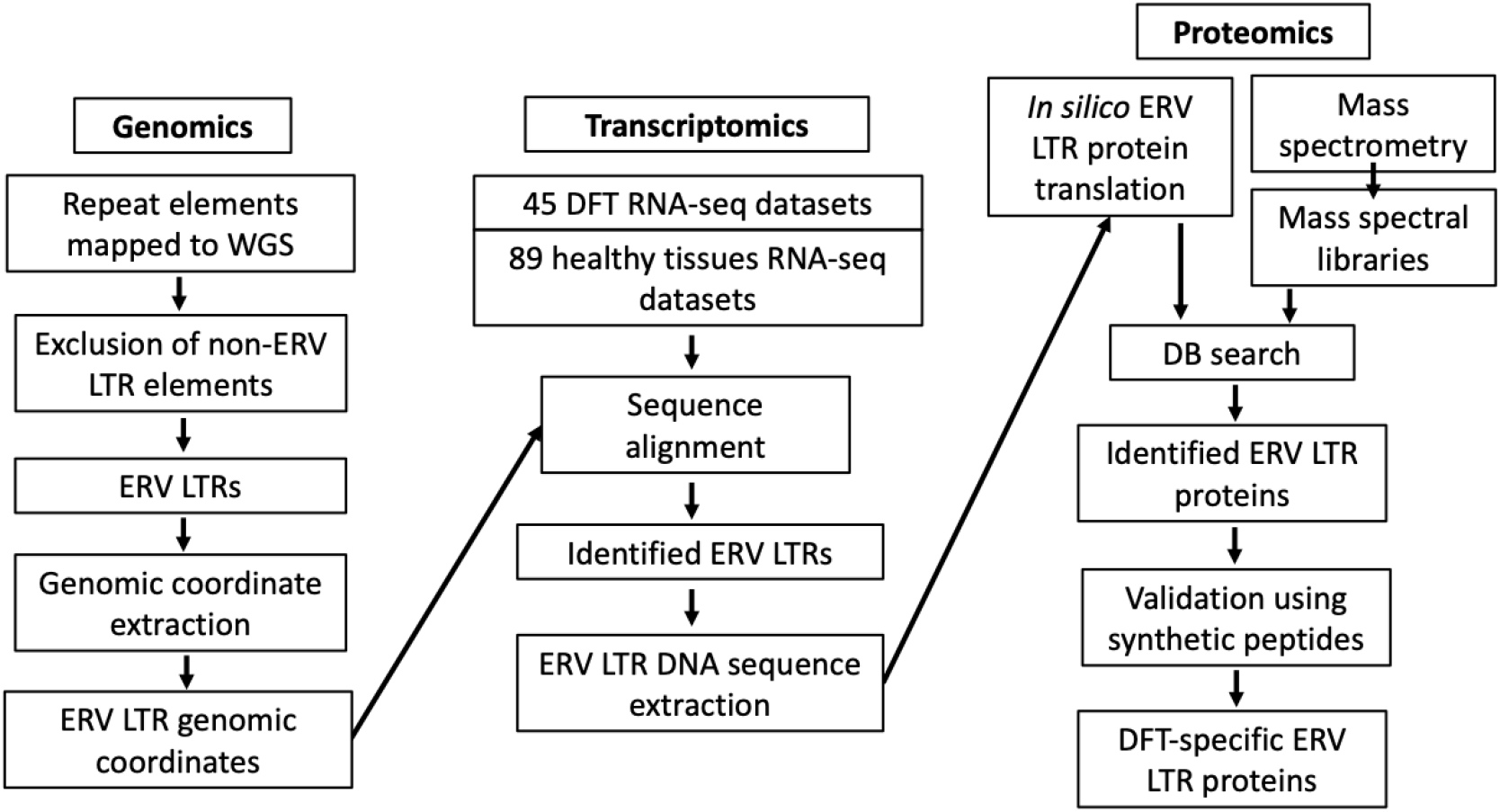
Overview of the ERV LTR proteogenomic pipeline. The genomics pipeline involves the mapping of repeat elements and extraction of the ERV LTR genomic coordinates. For the transcriptomics component, the coordinates from the previous step were used to identify ERV LTR mRNA transcripts uniquely expressed in DFT1 and/or DFT2 cells and tissues. In the proteomics pipeline, the coding sequence of these mRNA were predicted and used to identify ERV LTR peptides exclusively found in DFT1 and DFT2 immunopeptidomes.

### Mapping ERV LTRs in the devil genome

The RepeatMasker software [23] was used to identify ERV LTRs on the Tasmanian devil reference genome, mSarHar1.11 [4]. The dataset was generated using RepeatMasker v4.0.8 in sensitive mode, with Blastp version 2.0MP-WashU, utilising a combined database consisting of Dfam_Consensus-20181026 and RepBase-20181026. The following parameters were used: RepeatMasker -engine wublast -species ‘sarcophilus harrisii’ -s -no_is -cutoff 255 -frag 20000. Genomic coordinates for these candidates including the strand orientation were then extracted. The list of ERV LTRs was then combined with the list of canonical genes (Ensembl mSarHar1.11.110.gtf) to generate a custom ERV LTRs – normal genes database.

### Transcriptome data

See supplementary **Table S1** for the complete sample and RNA-seq dataset list. Nine DFT samples and twelve healthy samples were collected as fresh samples; all other transcriptome data were obtained from public data repositories [3, 5, 24, 25].

Eight primary DFT1, one DFT2, and two ear fibroblast biopsies were taken from wild devils using a 4 mm disposable biopsy punch (Kai Medical, #EUR0450/10) in accordance with the University of Tasmania Animal Ethics (permit IDs A0011436, A0012513, A0013685, A0014976, and A0014599). Biopsies were stored in RNAlater (Qiagen) except for fibroblast samples which were stored in transport medium consisting of RPMI medium (GIBCO) supplemented with 1% Antibiotic-Antimycotic (GIBCO). Single cell suspensions were then obtained from fibroblast samples by mincing. Cell suspension was cultured in RPMI medium supplemented with 1% Antibiotic-Antimycotic and 10% foetal calf serum, at 35 °C and 5% CO2 in a humidified incubator until ready to be harvested for RNA extraction. Two primary blood vessel samples were collected from euthanised devils without DFTD due to other animal welfare concerns, and stored in RNAlater. Whole-blood samples were collected from the jugular vein of wild healthy animals and stored in lithium heparin anticoagulant tubes (Greiner Bio-One).

For RNA extraction, solid biopsies and fibroblast cell lines were extracted using NucleoSpin™ RNA, Mini kit (Macherey-Nagel) according to the manufacturer’s protocol. Whole-blood samples were extracted using QIAamp RNA Blood Mini Kit (Qiagen) according to the manufacturer’s protocol. RNA integrity was assessed using a Eukaryotic RNA 6000 Nano Kit and 2100 Bioanalyzer (Agilent). 100 base-pair single-end and pair-end mRNA sequencing for each sample was performed on the Illumina Hiseq-2000 and Illumina Hiseq-2500 platform (Illumina) according to the manufacturer’s guideline (**Table S1**). The sequencing quality of was assessed using FastQC (https://www.bioinformatics.babraham.ac.uk/projects/fastqc/) [26]. Remaining sequencing adaptors were trimmed using ILLUMINACLIP function of Trimmomatic [27]. Seed mismatches, palindrome clip threshold, and simple clip threshold were set to 2, 30, and 10, respectively. These datasets can be accessed via the European Nucleotide Archive repository (ENA) project number PRJEB98267.

### RNA sequencing data analysis

All RNA-seq datasets were aligned to the Tasmanian devil whole genome assembly mSarHar1.11 using STAR aligner according to the software manual [28]. Technical replicates were combined. This created coordinate sorted .bam files which were then fed into Subread featureCounts [29] to identify RNA-seq reads mapped to normal genes and ERV LTRs. Read counts were normalised using EDAseq [30] with the non-linear full quantile normalisation method and converted into transcripts per kilobase million (TPM) to account for varying gene lengths. Multi-dimensional scaling (MDS) plots were generated to assess similarities across all samples using the *plotMDS* function in the limma R package [31]. Box plots were generated to visualise the number of ERV LTRs across all samples and DFT-specific LTRs using ggplot2 (https://ggplot2.tidyverse.org/) in R [32]. Kruskal-Wallis test and Welch two sample t-test were performed in R.

Differential expression analysis was carried out using the variance modelling at the observational level (*voom*) [33] function from limma [31]. Then, linear modelling was carried out using *lmfit* and *contrast*.*fit* functions prior to calculating empirical Bayes moderation with *eBayes* function of limma [31]. Lastly, the *treat* function of limma [31] was used to calculate *p*-values from empirical Bayes moderated t-statistics with log-fold-change (logFC) greater or less than 1. Venn diagram was generated to show the number of genes differentially expressed (DE), *p*-value < 0.05, between DFT1 vs normal tissues, DFT2 vs normal tissues, and in both comparisons using *vennDiagram* function in limma [31]. The EnhancedVolcano package (https://github.com/kevinblighe/EnhancedVolcano) in R was used to generate volcano plots to visualise DE genes in DFT1 vs normal tissues and DFT2 vs normal tissue. The Heatmaps for the top 30 upregulated ERV LTRs in DFT1 vs normal tissues and DFT2 vs normal tissues (adjusted p-value < 0.05) were generated using ComplexHeatmap R package [34].

### ERV LTR sequence extraction and translation

The DNA sequences of ERV LTR transcripts with TPM = 0 in all normal tissues and TPM > 0 in DFT1 or DFT2 samples were extracted from the devil WGS using Samtools faidx package [35]. Due to increasing evidence of non-canonical start codons and reading frames [36-38], we decided to not only include open reading frames (ORFs) from ATG start codon to stop codons, but also between stop codons. Sequences between stop codons that are ≥ 21 nucleotides long were extracted and 6-frame translated using orfipy [39]. This generated ERV LTR peptide sequences that are at least 7 amino acids long.

### Analysis of immunopeptidomes

The immunopeptidome dataset was retrieved from the PRIDE repository PXD020614 (https://www.ebi.ac.uk/pride/archive/projects/PXD020614) [40]. Briefly, the immunopeptidomes from the public dataset were generated using DFT1 cell line 4906, DFT2 cell line RV, and a devil fibroblast control cell line SALEM. DFT1 cell line 4906 was stimulated with recombinant devil interferon gamma (IFN-g) for 17 hours to upregulate the expression of MHC-I on DFT1 cells. This was to upregulate MHC-I, which is not constitutively expressed on DFT1 cells. Approximately 10^9^ cells from each of the four experimental replicates for each cell line (DFT1+IFN-g, DFT2, and fibroblast cells) were used for peptide isolation [40]. Liquid chromatography-tandem mass spectrometry (LC-MS/MS) was used to detect acquire spectra. Full-scan MS spectra were acquired using an Orbitrap of Q-Exactive Plus mass spectrometer in data-dependent acquisition (DDA) mode (Thermo Fisher Scientific, Bremen, Germany), full details provided in [40].

A custom database containing the Tasmanian devil reference proteome (https://www.uniprot.org/proteomes/UP000007648) and 6-frame translated DFT-specific ERV LTR sequences was used as a search database to identify ERV peptides in the immunopeptidomes. PEAKS® Studio 11.5 build 20231206 (Bioinformatics Solutions Inc., Waterloo, ON, Canada) was used. The parent mass error tolerance was set to 20 ppm using monoisotopic mass with fragment mass error tolerance of 0.02 Daltons. The enzyme and digest mode were set to none and unspecific, respectively. Deamidation (NQ), oxidation (M), and cysteinylation were chosen as variable post translational modifications (PTMs). Peptide false discovery rate (FDR) was set to 5%, and FDR estimation with decoy-fusion function was selected.

### LC-MS/MS and data analysis for synthetic peptides

The top 11 ERV LTR peptides detected from immunopeptidomes using PEAKS were selected for validation using synthetic peptides. Candidates were chosen based on the peptide abundance or the confidence score (-10logP). They comprised of five DFT1-specific peptides, five DFT2-specific peptides, and one shared peptide between DFT1 and DFT2. Additionally, nerve growth factor receptor (NGFR) (known to be upregulated in DFT1 and DFT2) was used as a positive control. Peptides were synthesised by GenScript with the purity of ≥ 95% (**Table S2**).

The synthetic peptides were analysed using a one-hour DDA-MS method with the starting condition of 98% mobile phase A (water + 0.01% formic acid) and 2% mobile phase B (80/20 acetonitrile/water + 0.08% formic acid). This was followed by the liquid chromatography (LC) gradient listed in **Table 1**. The samples were injected using an LC ‘trap/elute’ configuration in which peptides were loaded onto a 20 mm x 75 µm PepMap 100 C18 trapping column at 5 µL/min then separated using a 250 mm x 75 µm PepMap 100 C18 analytical column at 300 nL/min. The same custom search database listed in the immunopeptidomics section was used. PEAKS® Studio 11.5 build 20231206 (Bioinformatics Solutions Inc., Waterloo, ON, Canada) was again used to analysed acquired DDA raw files with the same parameters described earlier. MSMS spectra were compared and plotted using Python 3.9.16 and Matplotlib 3.7.0 packages [41].

**Table 1.** Liquid chromatography gradient for mass spectrometric analysis.

| *Time (start – end, minute)* | *Buffer B (%)* |
| --- | --- |
| 9 - 35 | 10 - 25 |
| 35 - 40 | 25 - 45 |
| 41 - 46 | 95 |
| 46 - 60 | 2 |

## Results

### Identification of repeat elements in the Tasmanian devil genome and transcriptomes

We identified more than 6.5 Million repeat elements in the most recent assembly of the Tasmanian devil reference genome (mSarHar1.11) [4] (**Table S2**). These elements belong to diverse repeat classes/families, such as long interspersed nuclear elements (LINEs), short interspersed nuclear elements (SINES), ERV LTRs, and simple repeats. We excluded all non-ERV LTR elements from the list to be left with a total of 79,309 candidate ERV LTRs in the Tasmanian devil genome (**Table S2**).

For gene expression analysis, the MDS plot indicated that the majority of DFT1 and DFT2 samples cluster together closely but have distinct patterns of gene expression compared to all normal tissues when assessing the whole transcriptome, which includes LTRs (**Figure 3*a***). Normal tissues were grouped into healthy lip, neuronal normal, and non-neuronal normal (excluding normal lip) samples. The segregation of DFT1 and DFT2 samples away from other normal tissues was more pronounced when only assessing the expression of LTRs (**Figure 3*b***). The median expression of over 2,500 LTRs were identified in DFTs and normal tissue samples (**Figure 3*c***). DFT1 was found to express fewer LTRs compared to neuronal normal tissues, *p =* 0.0117 (Kruskal-Wallis tests), but more LTRs compared to non-neuronal normal tissues, *p =* 0.0409 (Kruskal-Wallis tests, **Figure 3*c***). DFT1 was not statistically different in LTR abundance compared to DFT2 or lip samples (Kruskal-Wallis tests, **Figure 3*c***). Neuronal normal samples had more expressed LTRs compared to lip and non-neuronal tissue samples, *=* 4.8e-05 and *p =* 0.0004, respectively (Kruskal-Wallis tests, **Figure 3*c***). Interestingly, no significant differences were found between DFT2 LTRs and all other samples (Kruskal-Wallis tests, **Figure 3*c***).

**Figure 3.**
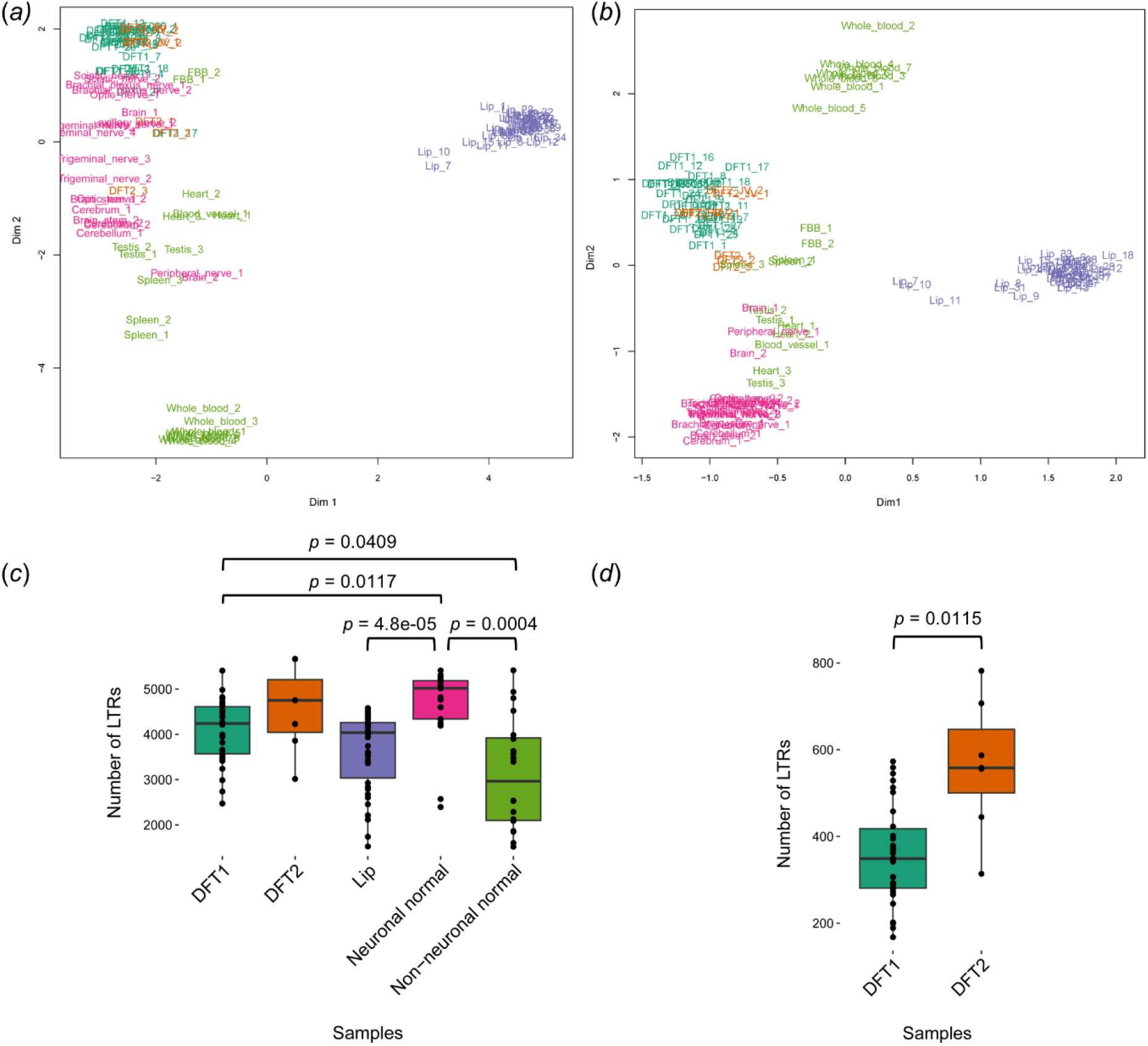
Multidimensional scaling (MDS) analysis of gene and LTR expression. MDS analysis showing similarity and dissimilarity of biological replicates across all sample types when assessing the gene expression of the whole transcriptome (including LTRs) (*a*) and LTRs independently (*b*). Dimensions (dim) 1 and 2 are shown on the X and Y axis, respectively. The average normalised number of LTRs identified in each sample (c). The error bars indicate the standard deviation (*c*). The average normalised number of LTRs exclusively identified in DFT1 and/or DFT2 samples (*d*). The error bars indicate the standard deviation. The p-values were calculated using Kruskal-Wallis tests in (*c*) and a Welch two sample t-test in (*d*). Sample types were colour coded as followed: DFT1, green; DFT2, orange; normal lip samples, purple; neuronal samples, pink; non-neuronal normal samples (excluding healthy lip samples), lime green.

When only looking at LTRs unique to DFT1 and/or DFT2, DFT1 samples had the median expression of slightly over 300 DFT-specific LTRs (Welch two sample t-test, **Figure 3*d***). Conversely, DFT2 had the median expression of marginally over 500 DFT-specific LTRs (Welch two sample t-test, **Figure 3*d***). Although DFT1 was not found to express a significant difference in total LTR numbers compared to DFT2 (Welch two sample t-test, **Figure 3*d***), it was found to express fewer DFT-specific LTRs in comparison to DFT2, *p =* 0.0115 (Welch two sample t-test, **Figure 3*d***).

### Differential expression analysis

Altogether, we identify a total of 4,962 differentially expressed genes including LTRs (**Figure S1**). **Figure 4*a*** shows differentially expressed LTRs between DFT1 vs normal, DFT2 vs normal, and common differentially expressed LTRs in DFT1 and DFT2. DFT1 and DFT2 had similar total numbers of differently expressed LTRs (**Figure 4*a***). The LTR expression landscapes of DFT1 or DFT2 compared to normal tissues are shown in **Figure 4*b*** and **4*c***, respectively. Additionally, genes that are known to be upregulated in DFT1 and DFT2, such as periaxin (PRX) in DFT1 [42, 43] and nerve growth factor receptor (NGFR) in DFT1 and DFT2 [5, 42] are shown to be differentially expressed in our study (**Figure S2**). The top 30 upregulated LTRs for DFT1 or DFT2 relative to normal tissues (adjusted p-value < 0.05, empirical Bayes moderated t-statistics) are shown in **Figure 5*a*** and **5*b***, respectively. These candidates were ranked based on the relative fold change (log2FC) compared to all normal tissues. Four of the top 30 upregulated LTRs are shared between DFT1 and DFT2 with log2FC > 4 (**Table 2**).

**Table 2.** Shared upregulated LTRs in DFT1 and DFT2 relative to normal samples.

| LTRS | DFT1 VS NORMAL |  | DFT2 VS NORMAL |  |
| --- | --- | --- | --- | --- |
|  | Log2FC | Adjusted p-value | Log2FC | Adjusted p-value |
| **25034_LTR200_MD** | 6.66 | 2.59E-35 | 6.65 | 7.60E-16 |
| **25033_LTR200_MD** | 5.78 | 8.11E-36 | 5.64 | 1.04E-13 |
| **66220_LTR200_MD** | 5.55 | 6.62E-16 | 5.94 | 6.95E-08 |
| **55447_ERV2-1_SH-I** | 4.77 | 2.19E-16 | 6.32 | 2.30E-11 |

**Figure 4.**
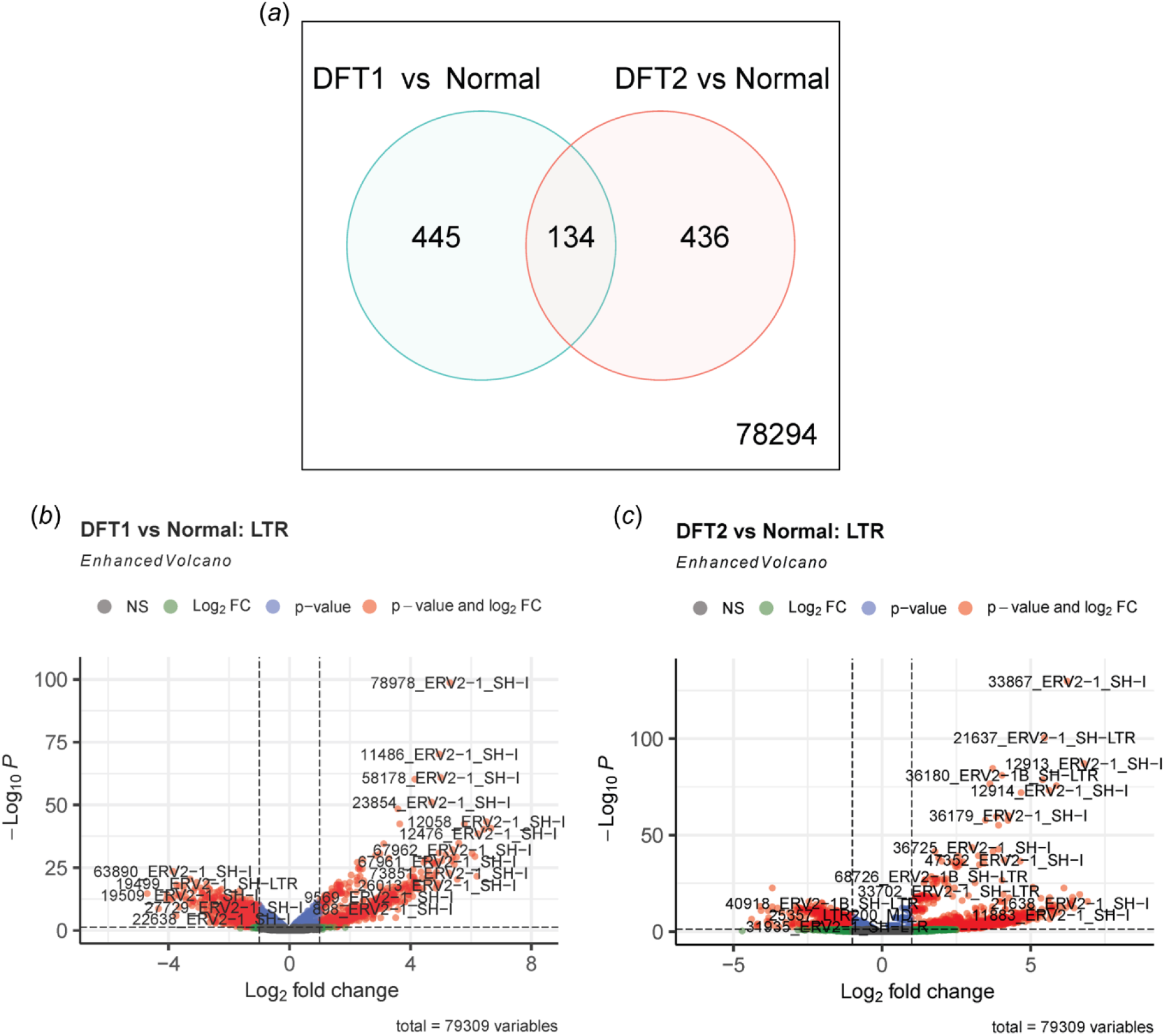
Venn diagram and volcano plots showing differentially expressed LTRs between DFT1 vs normal tissues and DFT2 vs normal tissues. Venn diagram showing differentially expressed LTRs between DFT1 vs normal tissues, DFT2 vs normal tissues, and common differentially expressed LTRs between DFT1 and DFT2 (*a*). Volcano plot showing differentially expressed LTRs of DFT1 vs normal tissues (*b*). Volcano plot showing differentially expressed LTRs of DFT2 vs normal tissues (*c*). Horizontal dotted line shows the p-value cut-off for 0.05 whilst vertical dotted lines show cut-off for |log2FC| of 1. NS denotes LTRs that are not differentially expressed.

**Figure 5.**
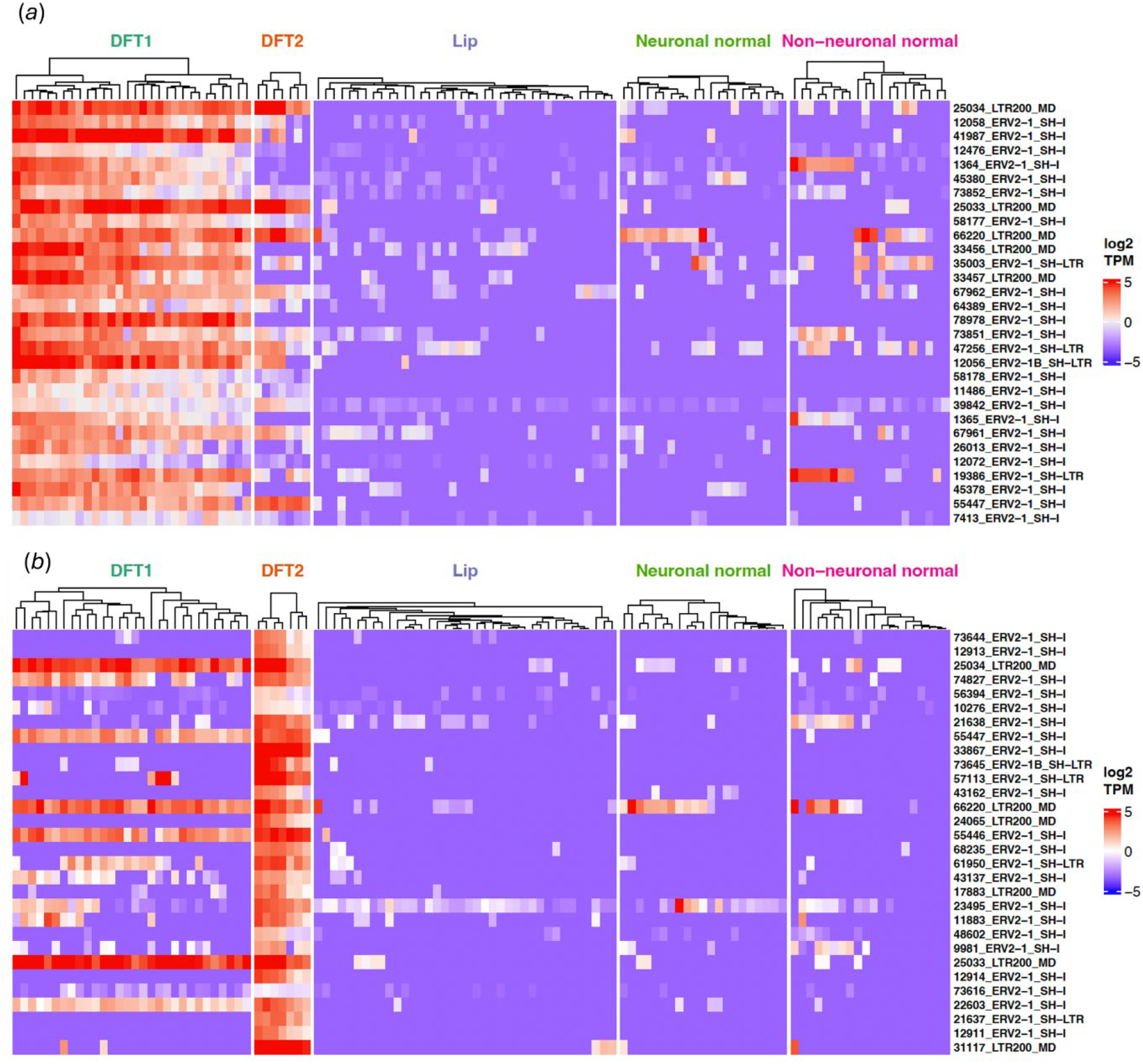
Heatmaps showing top 30 upregulated LTRs in DFT1 or DFT2 samples relative to normal tissues. Heatmap of the 30 most differentially expressed LTRs in DFT1 (*a*) and DFT2 (*b*) samples relative to normal tissues including lip, neuronal normal, and non-neuronal tissues. Dendrograms show the clustering of biological replicates within each tissue type (adjusted p-value < 0.05).

### Expression of LTR peptides in the immunopeptidome of DFT1 and DFT2 cells

Our study aimed to identify LTR peptides that are DFT1 and DFT2 specific, therefore, we only included those with no RNA transcripts detected in any of healthy samples (TPM = 0). These sequences were appended to the reference proteome to create a custom search database. Our analysis detected a total of 41,819 peptides across all samples (**Table S3**). Notably, our pipeline identified the same highly-expressed peptide (LHSDSGISVDSQS) derived from NGFR as previously reported [40]. We detected 69 unique LTR peptides across DFT1, DFT2, and fibroblast samples (**Figure 6*a*** and **Table S3**). Thirty-three LTR peptides were found only in DFT1 and DFT2 samples. Twenty-four LTR peptides were unique to DFT1 and eight LTR peptides were unique to DFT2 (**Figure 6*a***). One peptide was found in both DFT1 and DFT2 samples (LQDITQKI) (**Figure 6*a*** and **6*b***). Log2 converted relative abundances of LTR peptides identified in each sample is shown in **Figure 6*b***.

**Figure 6.**
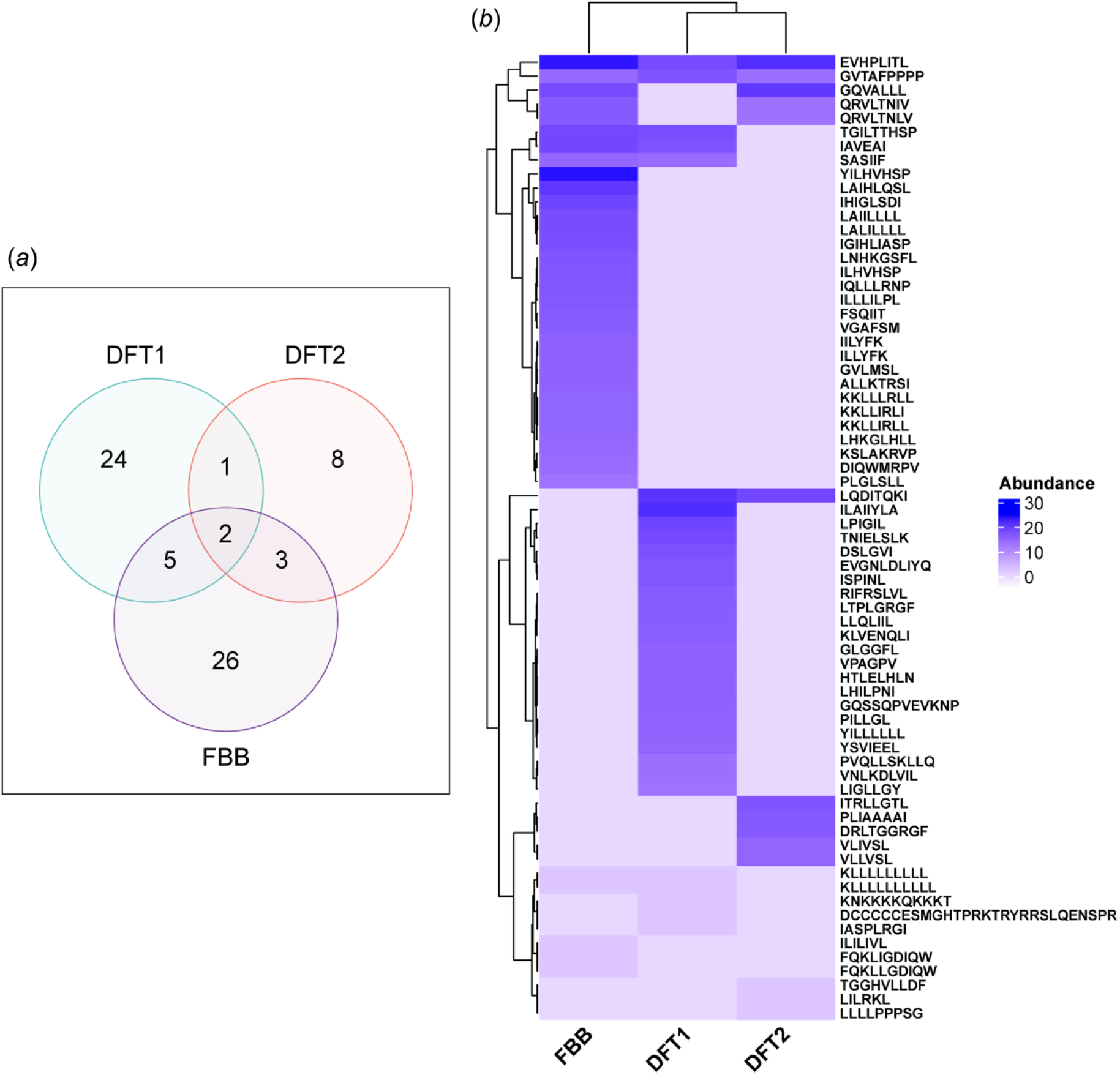
Venn diagram and heatmap showing identified LTR peptides. A Venn diagram showing LTR peptides identified across DFT1, DFT2, and fibroblast (FBB) samples (FDR < 0.05) (*a*). Heatmap displaying LTR peptides identified across DFT1, DFT2, and FBB samples (FDR < 0.05) (*b*). Peptide abundance was log2 converted from raw mass spectrometry relative ion intensity.

### Validation of LTR peptides with synthetic peptides

To confirm the correct assignment of ERV LTR peptides, we compared mass spectra of 11 LTRs identified by the database search with their synthetic peptide counterparts (**Table S3**). The selection criteria were based on the confidence score (-10logP as calculated by PEAKS) and peptide abundance. Additionally, NGFR peptide LHSDSGISVDSQS was included as a positive control. Mirror plots with Pearson correlation coefficients for common ions between experimental and synthetic peptide spectra were generated (**Figure 7**). This resulted in the detection of five LTR peptides - which consist of DRLTGGRGF, EVGNLDLIYQ, LQDITQKI, LPIGIL, and PLIAAAAI - based on the spectra matching and correlation score (**Figure 7**). These peptides were found in DFT1 or DFT2, except for LQDITQKI which was identified in both DFT1 and DFT2. Mirror plots of the LTR peptides predicted by the proteogenomic pipeline and the synthetic peptides that did not match are listed in **Figure S3**.

**Figure 7.**
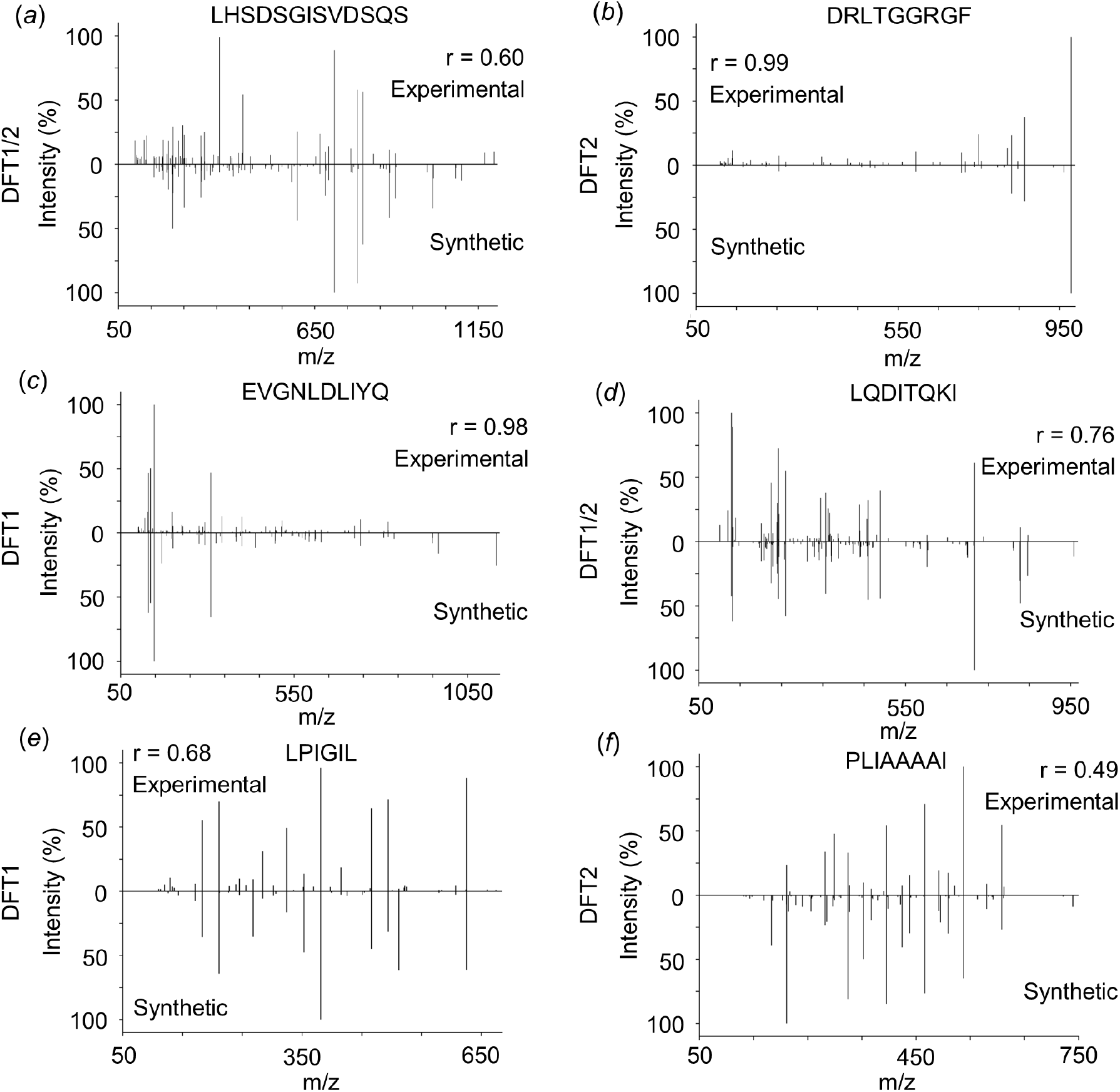
Mirror plots displaying mass spectra and Pearson correlation coefficient between pipeline predicted and synthetic peptides. NGFR peptide served as a positive control shown in (*a*) and ERV LTR peptides shown in (*b - f*). The DFT type where the peptide was identified is also shown. The x-axis shows the mass-to-charge (m/z) and y-axis shows the mass spectrometry intensity as percentage relative to the most intense peak. The upper half of the plot shows the mass spectra of the experimental peptides obtained from immunopeptidomes. The bottom half of the plot shows the mass spectra of the synthetic peptides. *r*, Pearson correlation coefficient.

## Discussion

Our proteogenomic analysis using a custom LTR detection pipeline identified several LTRs in the Tasmanian devil genome, and a subset of them were expressed in transcriptomes and immunopeptidomes. We identified 79,309 ERV LTRs in the devil’s genome, which was a similar number to a previous study conducted to investigate the presence of ERVs in dasyurids including Tasmanian devils [22]. This suggests that our pipeline is capable of accurately detecting and reporting LTRs found in the devil’s genome and transcriptome. Importantly, these DFT-specific LTRs found in immunopeptidomes may serve as potential targets for DFTD vaccines in development [6, 7].

Gene expression analysis showed that most LTR transcripts are not unique to DFT1 and DFT2, and thousands of LTRs are expressed across DFT, normal neuronal and non-neuronal samples. Hundreds of LTRs were exclusively identified in DFT1 and/or DFT2 transcriptomes. DFT2 expressed more LTRs than DFT1. Future studies should aim to include more DFT2 samples to improve the sample size balance between DFT1 and DFT2. Differential expression analysis revealed that some LTRs had significantly higher expression in DFT1 and/or DFT2 compared to all other normal tissues (Log2FC > 5). Four LTRs were found to be among the top 30 upregulated transcripts in both DFT1 and DFT2 relative to normal tissues. These shared LTRs may serve as potential therapeutic targets as tumour-associated antigens. As the scope for this study was to identify non-mutational TSAs derived from LTRs, we did not include LTRs transcripts detected in non-tumour transcriptomes in the immunopeptidomic analysis. However, future studies could assess the potential function roles of LTRs transcribed in non-tumour tissues.

Our analysis detected LTR peptides that were present in DFT1 and DFT2 but not present in the fibroblast immunopeptidome. Detected peptides from immunopeptidomes are particularly useful for the context of therapeutics development, as they demonstrate translation of LTRs and surface presentation on MHC-I complexes for cytotoxic T cell recognition. The recognition of foreign/non-canonical peptides can lead to an immune activation [44, 45]. We identified one LTR peptide that was shared between the two tumour types (LQDITQKI), making it an ideal vaccine target that can protect against both DFT1 and DFT2. Importantly, synthetic peptides corresponding to experimental peptides yielded similar spectra with moderate to high correlation score for 5 out of 11 LTR peptides chosen for mass spectrometric validation. This suggests that these are likely valid targets, not a result of misidentification.

A previous study reported that devil’s MHC-I preferentially binds to peptides that were 7-15 amino acid long (7-15mers) [40]. Our study identified a few 6mers, including LPIGIL which was validated with a synthetic peptide. Future studies should aim to validate whether smaller peptides can efficiently bind to the MHC-I molecule. Recombinant MHC-I refolding assay can be used to assess the biding affinity [46, 47]. Additional immunopeptidome datasets are needed to further explore the specificity of peptides to DFT1 and DFT2 cells. Furthermore, mapping of peptides to specific MHC-I alleles could provide insight into the potential for these ERV peptides to be expressed in devils with MHC-I alleles that match DFT1 or DFT2 cells. In humans, it was revealed that a subtype of human ERV is a critical regulator of Schwann cell proliferation in Merlin-Negative Schwannoma [48]. However, the links between ERVs or LTRs and DFT1/2 establishment and progression are yet unknown and require further investigation. The immunogenicity of these validated DFT-specific LTR peptides should be investigated to determine whether they can elicit an anti-tumour response that confers long term protection against DFTD in devils.

## Supporting information

Supplementary Table S1 List of RNA-seq datasets

Supplementary Table S2 Repeat elements identified in Tasmanian devils genome

Supplementary Table S3 List of identified peptides and synthetic peptides

## Acknowledgements

Thank you to the field teams of the Save the Tasmanian Devil Program and Rodrigo Hamede, Ruth J. Pye, and Camila Espejo who collected samples over the years to build the transcriptomic databases, and particularly Rodrigo Hamede for the DFT2 samples used in this study. We thank Jocelyn Darby for her contributions to the team. Thank you to Liz Murchison the genomics analysis that provided the foundation for this study.

## Funding

Australian Research Council grants LP210301148 (A.S.F, R.J.P), LP230200956 (A.S.F, R.J.P, K.A.F, R.W) and FT240100092 (A.S.F)

Wildcare Tasmania Nature Conservation Fund (A.S.F, R.J.P)

Federal Group through funds from Saffire Freycinet

University of Tasmania Advancement Office through funds raised by the Save the Tasmanian Devil Appeal (A.S.F, R.J.P), including support from the AAT Kings - Treadright Foundation, Smitten, and Pure Foods Eggs

Charitable organisation from the Principality of Liechtenstein (A.S.F)

Select Foundation Research Fellowship (A.S.F)

Tall Foundation (A.S.F)

## Author contributions

Field sample collection: R.J.P, C.E.

Laboratory analysis and bioinformatics: A.K., M.S., R.W., S.H.R, C.E.B.O., A.L.P., K.A.F.

Data analysis: A.K., M.S., C.E.B.O., A.L.P., K.A.F.

Funding acquisition; A.S.F, K.A.F.

Writing – original draft: A.K.

Writing – review and editing: A.K., M.S., R.W., S.H.R., C.E.B.O., K.A.F., A.B.L., A.L.P., A.S.F.

## References

1. Cunningham, C.X., et al., Quantifying 25 years of disease-caused declines in Tasmanian devil populations: host density drives spatial pathogen spread. Ecology Letters, 2021. 24(5): p. 958–969.

2. Pearse, A.M. and K. Swift, Allograft theory: transmission of devil facial-tumour disease. Nature, 2006. 439(7076): p. 549.

3. Pye, R.J., et al., A second transmissible cancer in Tasmanian devils. Proceedings of the National Academy of Sciences, 2016. 113(2): p. 374.

4. Stammnitz, M.R., et al., The evolution of two transmissible cancers in Tasmanian devils. Science, 2023. 380(6642): p. 283–293.

5. Patchett, A.L., et al., Two of a kind: transmissible Schwann cell cancers in the endangered Tasmanian devil (Sarcophilus harrisii). Cell Mol Life Sci, 2019.

6. Flies, A.S., et al., An oral bait vaccination approach for the Tasmanian devil facial tumor diseases. Expert Review of Vaccines, 2020. 19(1): p. 1–10.

7. Kayigwe, A.N., et al., A human adenovirus encoding IFN-γ can transduce Tasmanian devil facial tumour cells and upregulate MHC-I. Journal of General Virology, 2022. 103(11).

8. Tovar, C., et al., Regression of devil facial tumour disease following immunotherapy in immunised Tasmanian devils. Scientific Reports, 2017. 7(1): p. 43827.

9. Haen, S.P., et al., Towards new horizons: characterization, classification and implications of the tumour antigenic repertoire. Nature Reviews Clinical Oncology, 2020. 17(10): p. 595–610.

10. Apavaloaei, A., et al., The Origin and Immune Recognition of Tumor-Specific Antigens. Cancers (Basel), 2020. 12(9).

11. Smith, C.C., et al., Alternative tumour-specific antigens. Nature Reviews Cancer, 2019. 19(8): p. 465–478.

12. Cai, Y., et al., MHC-I-presented non-canonical antigens expand the cancer immunotherapy targets in acute myeloid leukemia. Scientific Data, 2024. 11(1): p. 831.

13. Laumont, C.M., et al., Noncoding regions are the main source of targetable tumor-specific antigens. Science Translational Medicine, 2018. 10(470): p. eaau5516.

14. Gonzalez-Cao, M., et al., Human endogenous retroviruses and cancer. Cancer Biol Med, 2016. 13(4): p. 483–488.

15. Grandi, N. and E. Tramontano, Human Endogenous Retroviruses Are Ancient Acquired Elements Still Shaping Innate Immune Responses. Frontiers in Immunology, 2018. 9.

16. Geis, F.K. and S.P. Goff, Silencing and Transcriptional Regulation of Endogenous Retroviruses: An Overview. Viruses, 2020. 12(8).

17. Dopkins, N. and D.F. Nixon, Activation of human endogenous retroviruses and its physiological consequences. Nature Reviews Molecular Cell Biology, 2024. 25(3): p. 212–222.

18. Kitsou, K., P. Lagiou, and G. Magiorkinis, Human endogenous retroviruses in cancer: Oncogenesis mechanisms and clinical implications. Journal of Medical Virology, 2023. 95(1): p. e28350.

19. Attig, J., et al., LTR retroelement expansion of the human cancer transcriptome and immunopeptidome revealed by de novo transcript assembly. Genome Res, 2019. 29(10): p. 1578–1590.

20. Gallus, S., et al., Evolutionary histories of transposable elements in the genome of the largest living marsupial carnivore, the Tasmanian devil. Mol Biol Evol, 2015. 32(5): p. 1268–83.

21. Nilsson, M.A., The devil is in the details: Transposable element analysis of the Tasmanian devil genome. Mobile Genetic Elements, 2016. 6(1): p. e1119926.

22. Harding, E.F., et al., Invasion and amplification of endogenous retroviruses in Dasyuridae marsupial genomes. Molecular Biology and Evolution, 2024.

23. Smit, A., Hubley, R & Green, P, RepeatMasker. 2013–2015.

24. Wilson, R., et al., Transcriptome and proteome profiling reveals stress-induced expression signatures of imiquimod-treated Tasmanian devil facial tumor disease (DFTD) cells. Oncotarget, 2018. 9(22).

25. Ong, C.E.B., et al., NLRC5 regulates expression of MHC-I and provides a target for anti-tumor immunity in transmissible cancers. Journal of Cancer Research and Clinical Oncology, 2021. 147(7): p. 1973–1991.

26. Andrews, S., FastQC: A Quality Control Tool for High Throughput Sequence Data. 2010.

27. Bolger, A.M., M. Lohse, and B. Usadel, Trimmomatic: a flexible trimmer for Illumina sequence data. Bioinformatics, 2014. 30(15): p. 2114–2120.

28. Dobin, A., et al., STAR: ultrafast universal RNA-seq aligner. Bioinformatics, 2013. 29(1): p. 15–21.

29. Liao, Y., G.K. Smyth, and W. Shi, featureCounts: an efficient general purpose program for assigning sequence reads to genomic features. Bioinformatics, 2014. 30(7): p. 923–30.

30. Risso, D., et al., GC-Content Normalization for RNA-Seq Data. BMC Bioinformatics, 2011. 12(1): p. 480.

31. Ritchie, M.E., et al., limma powers differential expression analyses for RNA-sequencing and microarray studies. Nucleic Acids Research, 2015. 43(7): p. e47–e47.

32. Wickham, H., ggplot2. WIREs Computational Statistics, 2011. 3(2): p. 180–185.

33. Law, C.W., et al., voom: precision weights unlock linear model analysis tools for RNA-seq read counts. Genome Biology, 2014. 15(2): p. R29.

34. Gu, Z., R. Eils, and M. Schlesner, Complex heatmaps reveal patterns and correlations in multidimensional genomic data. Bioinformatics, 2016. 32(18): p. 2847–2849.

35. Li, H., A. Whitwham, and R. Davies. samtools faidx – indexes or queries regions from a fasta file. 2019; Available from: http://www.htslib.org/doc/samtools-faidx.htm.

36. Laumont, C.M., et al., Global proteogenomic analysis of human MHC class I-associated peptides derived from non-canonical reading frames. Nature Communications, 2016. 7(1): p. 10238.

37. Andreev, D.E., et al., Non-AUG translation initiation in mammals. Genome Biology, 2022. 23(1).

38. Kearse, M.G. and J.E. Wilusz, Non-AUG translation: a new start for protein synthesis in eukaryotes. Genes & Development, 2017. 31(17): p. 1717–1731.

39. Singh, U. and E.S. Wurtele, orfipy: a fast and flexible tool for extracting ORFs. Bioinformatics, 2021. 37(18): p. 3019–3020.

40. Gastaldello, A., et al., The immunopeptidomes of two transmissible cancers and their host have a common, dominant peptide motif. Immunology, 2021.

41. Hunter, J.D., Matplotlib: A 2D Graphics Environment. Computing in Science & Engineering, 2007. 9(3): p. 90–95.

42. Murchison, E.P., et al., The Tasmanian Devil Transcriptome Reveals Schwann Cell Origins of a Clonally Transmissible Cancer. Science, 2010. 327(5961): p. 84–87.

43. Tovar, C., et al., Tumor-Specific Diagnostic Marker for Transmissible Facial Tumors of Tasmanian Devils. Veterinary Pathology, 2011. 48(6): p. 1195–1203.

44. Seder, R.A. and R. Ahmed, Similarities and differences in CD4+ and CD8+ effector and memory T cell generation. Nature Immunology, 2003. 4(9): p. 835–842.

45. Rospo, G., et al., Non-canonical antigens are the largest fraction of peptides presented by MHC class I in mismatch repair deficient murine colorectal cancer. Genome Medicine, 2024. 16(1): p. 15.

46. Du, H., et al., Targeting peptide antigens using a multiallelic MHC I-binding system. Nature Biotechnology, 2024.

47. Rasmussen, M., et al., Uncovering the Peptide-Binding Specificities of HLA-C: A General Strategy To Determine the Specificity of Any MHC Class I Molecule. The Journal of Immunology, 2014. 193(10): p. 4790–4802.

48. Maze, E.A., et al., Human Endogenous Retrovirus Type K Promotes Proliferation and Confers Sensitivity to Antiretroviral Drugs in Merlin-Negative Schwannoma and Meningioma. Cancer Research, 2022. 82(2): p. 235–247.

