## Supplementary figures S1-S3 for "Identification of Expressed Endogenous Retroviral Element Associated Long Terminal Repeats in Devil Facial Tumour Cells"

GPO Box 341

Hobart, Tasmania, 7001, Australiat: +61 3 6226 4614

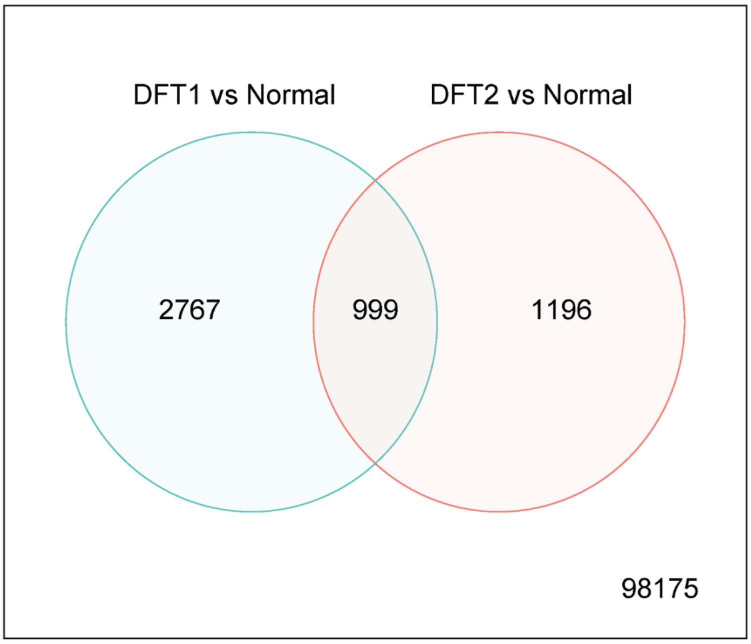


**Supplementary Figure S1. Venn diagram showing differently expressed genes between DFT1 vs normal tissues and DFT2 vs normal tissues.**

Venn diagram showing all differentially expressed genes, including LTRs, between DFT1 vs normal tissues, DFT2 vs normal tissues, and common differentially expressed genes between DFT1 and DFT2.

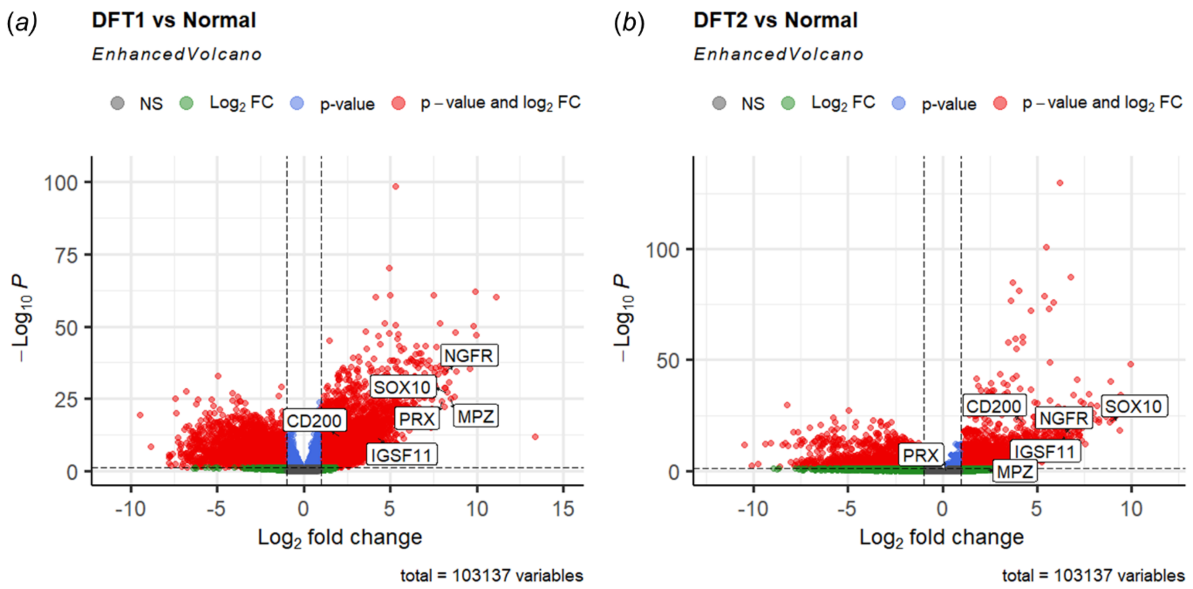


**Supplementary Figure S2. Volcano plots highlighting genes that are known to be upregulated in DFT1 and/or DFT2.**

Volcano plot showing differentially expressed LTRs of DFT1 vs normal tissues (*b*). Volcano plot showing differentially expressed LTRs of DFT2 vs normal tissues (*c*). Horizontal dotted line shows the p-value cut-off for 0.05 whilst vertical dotted lines show cut-off for |log2FC| of > 1. NS denotes LTRs that are not differentially expressed.

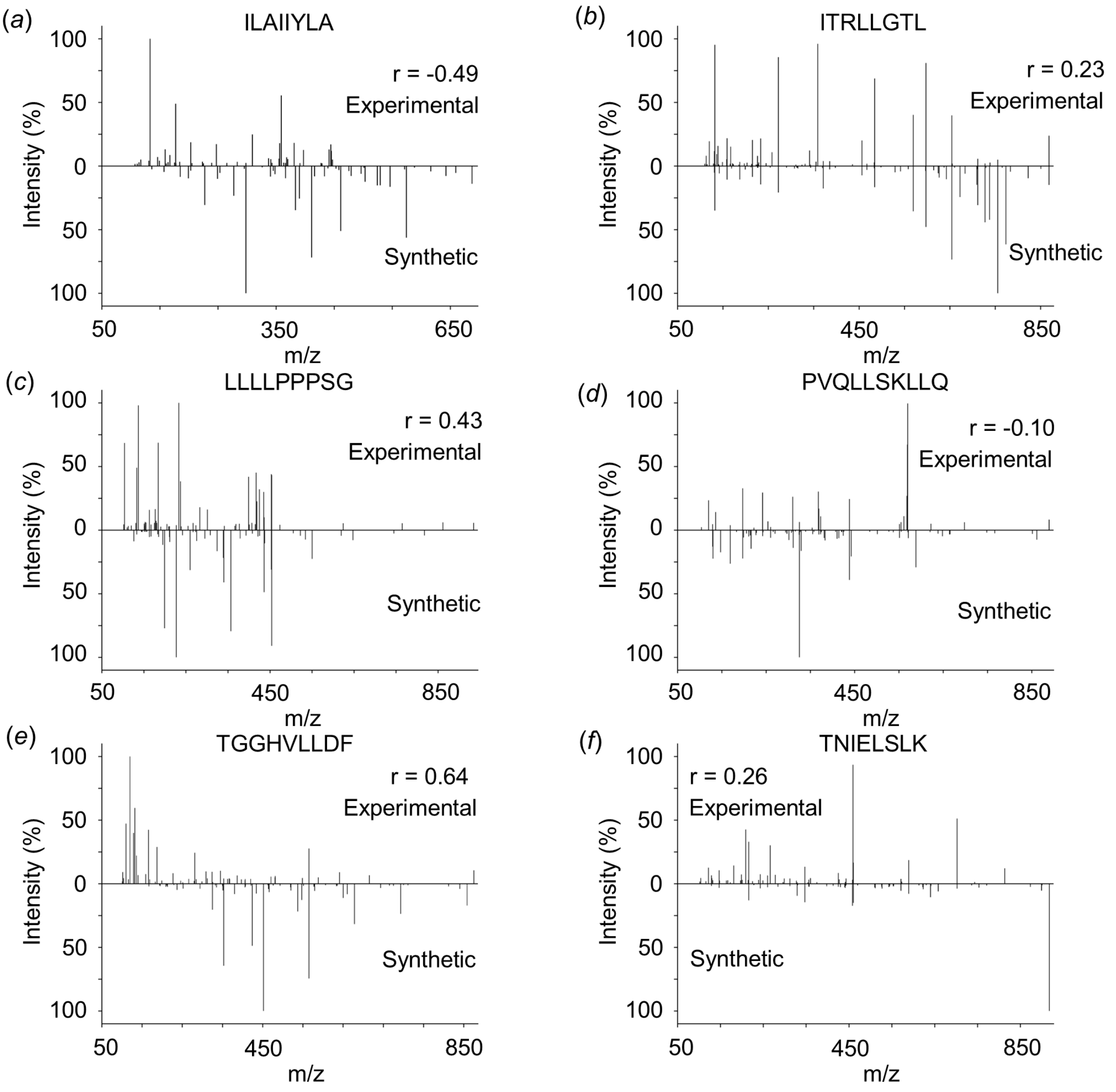


**Supplementary Figure S3. Mirror plots displaying mass spectra and Pearson correlation coefficient between native and synthetic peptides of likely misidentified canddates.** ERV LTR peptides are shown in (*a* - *f*). The x-axis shows the mass-to-charge (m/z) and y-axis shows the mass spectrometry intensity as percentage relative to the most intense peak. The upper half of the plot shows the mass spectra of the experimental peptides obtained from immunopeptidomes. The bottom half of the plot shows the mass spectra of the synthetic peptides. *r*; Pearson correlation coefficient.
