## Supplementary Table S2 Repeat elements identified in Tasmanian devils genome for "Identification of Expressed Endogenous Retroviral Element Associated Long Terminal Repeats in Devil Facial Tumour Cells"

GPO Box 341

Hobart, Tasmania, 7001, Australiat: +61 3 6226 4614

**Supplementary Table S2 Repeat elements identified in Tasmanian devils genome**

The dataset has been doposited on https://doi.org/10.5281/zenodo.17282419. The dataset contains a list of repeat elements identified in the Tasmanian devil (*Sarcophilus harrisii*) genome (mSarHar1.11). For further details, please see https://doi.org/10.1126/science.abq6453. The dataset was generated using RepeatMasker v4.0.8 in sensitive mode, with Blastp version 2.0MP-WashU, utilising a combined database consisting of Dfam_Consensus-20181026 and RepBase-20181026. The following parameters were used: RepeatMasker -engine wublast -species 'sarcophilus harrisii' -s -no_is -cutoff 255 -frag 20000.
